# Complete cross strain protection against congenital cytomegalovirus infection requires a vaccine encoding key antibody (gB) and T-cell (immediate early 1 protein) viral antigens

**DOI:** 10.1101/2025.06.18.660432

**Authors:** K. Yeon Choi, Yushu Qin, Nadia El-Hamdi, Alistair McGregor

## Abstract

**Background:** Cytomegalovirus is a leading cause of congenital disease and multiple strains enable congenital CMV (cCMV) from both primary and non-primary infection. A cross-strain protective cCMV vaccine is a high priority. The guinea pig is the only small animal model for cCMV and guinea pig cytomegalovirus (GPCMV) encodes functional homolog proteins including cell entry gB glycoprotein and non-structural immediate early 1 protein (IE1), essential for lytic infection. A gB vaccine antibody response fails to provide horizontal protection against highly cell-associated clinical GPCMV strain TAMYC compared to prototype strain 22122. Previously, a recombinant defective adenovirus (Ad) vaccine encoding IE1, a T-cell antigen, provided high-level cCMV protection. In this study, we hypothesized that a combined Ad-based strategy encoding trimeric gB complex and IE1 (AdgB+AdIE1) could improve cross-strain protection against cCMV compared to a gB vaccine (AdgB).

**Methods:** A preconception vaccine study evaluated the immune response and ability of vaccines to provide cross-strain protection against cCMV. Seronegative female animals were assigned into three vaccine groups: Group 1 (AdgB); Group 2 (AdgB+AdIE1); Group 3 (no vaccine). Animals were vaccinated following a previously defined protocol and antibody ELISAs were used to evaluate gB immune response (AD1, prefusion gB and wild type gB). Additionally, an IFNγ-ELISPOT assay evaluated IE1 T-cell response. During second trimester dams were challenged with GPCMV (22122 and TAMYC) and pregnancy went to term where viral loads in pup target organs and placentas were evaluated.

**Results:** Vaccinated dams elicited a higher neutralizing antibody response to gB than natural convalescent immunity and antibodies recognized homolog AD1 gB domain as well as prefusion gB with response surpassing natural immunity. Group 2 animals additionally elicited a T-cell response to IE1. Evaluation of viral load in pups demonstrated that AdgB+AdIE1 vaccine reduced GPCMV transmission to below detectable limits compared to 91.7% in unvaccinated group. In contrast, AdgB reduced cCMV transmission to 12% in pups.

**Conclusion:** Complete cross-strain cCMV protection is a significant milestone in this model and achieved by inclusion of an antibody response to trimeric gB and T-cell response to IE1. Importantly, gB and IE1 responses can synergize and increase protection against cCMV unlike prior approaches.

## 1 Introduction

Human cytomegalovirus (HCMV) is a ubiquitous pathogen establishing lifelong infection with disease in immunosuppressed populations including transplant and AIDS patients. Additionally, HCMV has the ability to cross the placenta and cause congenital cytomegalovirus (cCMV) infection with symptomatic disease causing vision and cognitive impairment as well as sensorineural hearing loss (SNHL) in newborns and is associated with autism (1, 2). In Europe and the US, cCMV occurs in approximately 0.5-1.2% of newborn babies with up to 30% of hearing loss in children attributed to cCMV (3–5). Globally over a million babies are born each year with cCMV (1) and a vaccine is considered a high priority (6). Perhaps the greatest risk of cCMV is to mothers who acquire a primary infection during pregnancy (7). In the US, it is estimated that 50% of women of childbearing age are CMV negative (8). However, a vaccine against cCMV is complicated by the existence of multiple strains of HCMV in the population and convalescent natural immunity is insufficient to prevent re- infection (7). Therefore cCMV can occur as a result of non-primary infection in women convalescent for the virus as a result of infection by a new strain and/or impaired immunity against the virus (1).

Importantly, the level of cCMV related to non-primary infection is an underappreciated burden, especially on a global scale or in countries or regions of high endemic CMV levels where cCMV levels can be as high as 5% (7, 8). Hence, the challenge of attaining a successful licensed vaccine against cCMV cannot be understated with a requirement to greatly exceed protective levels of natural convalescent immunity. Despite 50 years of vaccine research, a licensed cCMV vaccine remains an elusive goal and likely antibody and T-cell responses to key target antigens are required.

HCMV species specificity complicates studies in preclinical animal model, which requires the use of species-specific animal CMV. Additionally, there are only two animal models for cCMV (guinea pig and rhesus macaques). Currently, no vaccine strategy has been evaluated against cCMV in the rhesus macaque non-human primate (NHP) model. The guinea pig is the only small animal model for cCMV with guinea pig cytomegalovirus (GPCMV) causing disease in newborn pups similar to humans including SNHL (9–12). The guinea pig has a hemomonochorial placenta structure similar to human and a gestation period of approximately 70 days, which allows pregnancy studies to be evaluated in trimesters with prenatal pup neuro-anatomical development occurring almost completely in utero (13–15). Similarity of GPCMV and HCMV cell tropism, cell entry pathways and functional homolog proteins as vaccine target antigens, further cement the importance of this model for congenital and systemic disease studies (12, 16–22). The majority of GPCMV research to date has focused on prototype GPCMV 22122 strain. Despite the ability of 22122 virus to cause cCMV, a focus on this strain for vaccine protection studies does not accomplish an important objective for a translational CMV vaccine model i.e. cross-strain virus protection. A novel clinical strain of GPCMV (designated TAMYC) isolated from the salivary gland of a commercial animal was used by our laboratory to evaluate cross strain vaccine efficacy resulting in lower levels of protection in a non-pregnant animal model compared to studies with 22122 strain (12, 19–21, 23–25).

A significant focus of CMV vaccine studies is directed towards antibody response against viral glycoprotein complexes for cell entry (26–32). GPCMV has similar glycoprotein complexes to HCMV (gB, gH/gL/gO, gM/gN), which are essential for GPCMV direct pathway of infection of fibroblast cells as well as neutralizing target antigens (17, 20, 33–36) . Additionally, both HCMV and GPCMV encode a glycoprotein pentamer complex (PC) that is necessary for infection of non- fibroblast cells via an endocytic entry pathway (17, 22, 37, 38). In both HCMV and GPCMV, the gB glycoprotein is an immunodominant antibody target and essential for infection of all cells (17, 33, 39–41). A HCMV gB subunit vaccine, despite evoking high antibody titers, only provided approximately 50% efficacy in phase II clinical trials (42, 43). GPCMV gB subunit vaccine strategies in the guinea pig model fail to fully protect against cCMV and provides similar efficacy to that of human gB vaccine trials (44). However, the majority of GPCMV gB vaccine studies have only evaluated monomeric gB and a homotrimeric gB vaccine strategy has the capacity to improve neutralizing antibody response (20). Additionally, inclusion of the PC in a GPCMV vaccine improves virus neutralization on non-fibroblast cell lines and can enhance protection against cCMV (19, 34). However, this approach is less effective against highly cell associated GPCMV (TAMYC) clinical strain (24).

In HCMV, convalescent patients produce T cell responses against two specific viral proteins of importance: viral tegument protein pp65; and non-structural IE1 protein (45). GPCMV encodes functional homologs to IE1 and pp65 (GP83) involved in innate immune evasion (21, 46). Both antigens induce a cell-mediated immune response in guinea pigs (34, 47, 48). Various pp65 (GP83) vaccine strategies against cCMV in the guinea pig model had moderate levels of success and inclusion of both gB and pp65 in a vaccine approach did not improve cCMV vaccine efficacy against 22122 strain (47, 49–51). Additionally, the pp65 antigen provided poor cross strain protection against clinical strain GPCMV (TAMYC) despite 100% identity in pp65 amino acid sequence between strains (21). Recently, we demonstrated that GPCMV IE1 used in a recombinant Ad vaccine platform (AdIE1) provided high level cross strain protection against cCMV (48). In this report, we investigated the hypothesis that GPCMV cross strain vaccine efficacy against cCMV could be improved compared to prior AdIE1 vaccine by combining immunodominant antibody target (trimeric capable GPCMV gB glycoprotein) with IE1 T cell antigen in a recombinant Ad vaccine strategy (AdgB+AdIE1) in comparison to a gB (AdgB) only vaccine or no vaccine control study group. Novel GPCMV gB antibody ELISAs were developed for prefusion gB and a homolog AD1 gB domain that is immunodominant for gB in HCMV. Vaccine strategies induced a higher antibody response to gB compared to natural GPCMV convalescent immunity with neutralizing antibodies effective against both prototype (22122) and clinical (TAMYC) strains of GPCMV. Additionally, AdgB+AdIE1 vaccine induced a cell mediated response to IE1 unique protein sequence (GP123) in an IFNγ ELISPOT assay. In a preconception vaccine strategy, the AdgB+AdIE1 combined vaccine provided the first successful approach for complete cross strain protection against cCMV in this preclinical animal model. The AdgB+AdIE1 vaccine group had no detectable virus in pup litters compared to detectable virus in both the no vaccine control group and standalone AdgB vaccine group. Results demonstrate the combined importance of IE1 and gB as key antigens for a successful protective vaccine strategy against cCMV evoking both humoral (gB) and cell mediated (IE1) immune response against CMV.

## 2 Materials and Methods

### 2.1 Cells, viruses, oligonucleotides and genes

GPCMV (strain 22122, ATCC VR682) was propagated on various cell lines. Guinea pig fibroblast lung cells (GPL; ATCC CCL 158) and renal epithelial cells (REPI) for specific tropism studies as previously described (17, 18). Both clinical strain GPCMV TAMYC (25) and 22122 strain were expanded from separate guinea pig salivary gland stocks and exclusively propagated on REPI cells for maintenance of fully tropic virus stocks and animal challenge studies (17). Recombinant defective adenovirus (Ad5) vectors (E1 and E3 deleted) were previously described (20, 48) encoding full length gB (AdgB) or IE1 cDNA (AdIE1) with ORFs under HCMV IE enhancer control with 3’ SV40 polyA sequence. High titer CsCl gradient purified recombinant defective adenovirus virus stocks (10^12^ TDU/ml) were generated by Welgen Inc. (MA). All oligonucleotides were synthesized by Sigma-Genosys (The Woodlands, TX). Synthetic codon optimized genes were developed for: (1) prefusion gB (prefgB) and (2) gB AD1 domain as a fusion protein with gB(AD1) expressed at the C-terminal of GST, designated GST-gB(AD1). Based on alignment of GPCMV gB with HCMV gB protein amino acid sequences, a GPCMV prefgB was generated by gene synthesis (Genscript) incorporating removal of the furin site and additional specific amino acid substitutions equivalent to those made in HCMV gB ectodomain to generate a locked version of GPCMV prefgB (52). The ORF additionally incorporated a C-terminal FLAG epitope tag. The GPCMV prefgB was cloned into pcDNA3.1(+) vector (Invitrogen) under HCMV IE enhancer promoter control to enable transient expression in plasmid transfected cells (see ELISA section 2.5.1).

### 2.2 Ethics

Guinea pig (Hartley) animal studies were carried out under IACUC (Texas A&M University). All study procedures were carried out in strict accordance with the recommendations in the “Guide for the Care and Use of Laboratory Animals of the National Institutes of Health.” Animals were observed daily by trained animal care staff, and animals that required care were referred to the attending veterinarian for immediate care or euthanasia. Terminal euthanasia was carried out by lethal CO_2_ overdose followed by cervical dislocation in accordance with IACUC protocol and NIH guidelines. Animals purchased from Charles River Laboratories were verified as seronegative for GPCMV by toe nail clip bleed and anti-GPCMV ELISA of sera as previously described (33). Animal studies were carried out to determine: (1) GPCMV (TAMYC strain) pathogenicity in seronegative or seropositive (22122 strain) animals; (2) Immune response to natural convalescent immunity (GPCMV 22122 strain); (3) Immune response of candidate vaccines (AdgB or AdgB+AdIE1); (4) Vaccine efficacy against congenital GPCMV challenge (22122 and TAMYC strains) and protection of pups in vaccine group compared to no vaccine control group.

### 2.3 Animal studies

(A) Natural immunity and protection from GPCMV infection. GPCMV (22122) convalescent immunity in animals and cross strain protection against TAMYC strain challenge (SQ). Animal studies were performed to evaluate protection against TAMYC strain GPCMV challenge in animals convalescent for 22122 strain. Seronegative female guinea pigs (n=12) were injected once subcutaneously with the 22122 strain GPCMV (10^5^ pfu) at 3 months post challenge immune response was evaluated (anti-GPCMV) to verify seroconversion status and designated as seropositive animal group. Pooled sera were used for additional evaluation of immune responses to specific viral glycoprotein complexes (Supplementary Figure S1) and in virus neutralization studies. A second group of animals (n=12) were verified and used as seronegative animal group. Animals in both groups were challenged with TAMYC strain GPCMV (10^5^ pfu, SQ) at day 0. At 4, 8, 12 and 27 days post-infection (DPI), three animals per group were euthanized to evaluate viral load in target organs and blood via real-time PCR assay as previously described (23). (B) AdgB or AdgB+AdIE1 preconception vaccine protection against congenital GPCMV (TAMYC and 22122 strains). Seronegative female guinea pigs were randomly assigned to two different groups. Group 1 (AdgB vaccine group; n=17) or Group 2 (AdgB+AdIE1 vaccine group; n=13) vaccinated SQ with corresponding vaccine (10^8^ TDU) and boosted 4 and 8 weeks post original vaccine with equivalent dosage. Group 3 animals (n=10) were assigned as no vaccine control group. At 4 weeks post last vaccination, dams were paired with seronegative males for mating. The control group was similarly paired for mating. Dams were confirmed to be pregnant by palpation at approximately days 20 to 25 of gestation. At the late second trimester/early third trimester, pregnant animals in all groups were challenged with both strains of wild type GPCMV (22122 and TAMYC strains) in separate SQ injections (10^5^ pfu GPCMV/injection) into opposite flanks, and the animals were allowed to go to term. Subsequently, the viral load in the target organs (liver, lung, spleen, brain) of live-born or still- born pups was evaluated by real-time PCR. Placenta tissue when available was also evaluated for viral load.

### 2.4 Real time PCR assay

Tissues were collected from euthanized guinea pigs to determine the viral load. For tissue DNA extraction, FastPrep 24 (MP Biomedicals) was used to homogenize tissues as a 10% weight/volume homogenate in Lysing Matrix D (MP Biomedicals). DNA was extracted using the QIAcube HT (Qiagen) according to manufacturer’s tissue protocol instructions. Viral load was determined by real time PCR on a Lightcycler 480 (Roche Applied Science) as previously described (17, 34). Primers and hydrolysis probe were designed using the Lightcycler Probe Design2 program to amplify a product from the GPCMV viral polymerase subunit *GP44* gene: Forward primer 5’TCTCCACGGTGAAAGAGTTGT; Reverse primer 5’GTGCTGTCGGACCACGATA; hydrolysis probe 5’FAM-TCTTGCTCTGCAGGTGGACGA-BHQ1. Data was analyzed with the LightCycler Data Analysis Software (Version 1.5.1; Roche). Standard curve was generated using serial dilutions of GPCMV 22122 *GP44* plasmid at known concentrations for quantification and assay sensitivity.

The same standard curve was generated using TAMYC *GP44* plasmids to demonstrate that Universal GP44 primer-probe set can detect both strains. The sensitivity of the assay was determined to be 2 copies/reaction. Viral load was expressed as copy number/mg tissue or copies/ml of blood. Results calculated were a mean value of triplicate PCR runs per sample.

### 2.5 ELISA and Western blot assays

#### 2.5.1 ELISA

Anti-GPCMV ELISA was carried out as previously described (33) to determine GPCMV sero-status of commercially obtained guinea pigs (Charles River) as commercial animal colonies are not negative for GPCMV. Specific glycoprotein complex ELISAs (gB, gM/gN, gH/gL and PC) were carried out as previously described (19) using positive coating antigen derived from renal epithelial cell monolayers transduced with recombinant replication defective adenovirus (Ad) vectors expressing specific viral glycoprotein complexes or recombinant Ad vector expressing GFP for negative coating antigen (17, 33, 34). In the case of gM/gN, codon optimized synthetic genes in mammalian plasmid expression vectors were used instead (19, 33). Novel gB glycoprotein assays developed were for prefgB and gB(AD1) ELISAs. For prefgB, a synthetic prefusion gB mammalian expression plasmid (pcDNA3.1(+), Invitrogen) was used to express prefgB in REPI cells to generate coating antigen following a similar protocol for gM/gN ELISA (19, 33). Expression of prefgB was verified by Western blot (Supplementary Figure S2). A homolog AD1 domain was previously identified between HCMV gB and various animal CMV gB proteins based on BLAST alignment (53). An alignment of GPCMV gB and HCMV gB ORFs encoding AD1 region is shown in Supplementary Figure S3A. The minimal AD1 region was extended for GPCMV gB with additional flanking N and C terminal amino acids to match an extended AD1 sequence used to express a commercial HCMV gB(AD1) protein as a GST-gB(AD1) fusion protein (Sigma). A synthetic GST protein ORF encoding C-terminal fused GPCMV gB(AD1) region (gB amino acid codons 524-646) was cloned into a bacterial expression vector and expressed as a recombinant protein (Genscript) with recombinant protein purified by GST affinity chromatography for use in gB(AD1) ELISA studies. For gB(AD1) ELISA, a recombinant protein GST-gB(AD1) was expressed in bacteria and recombinant protein purified by GST affinity column chromatography (Genscript) and recombinant protein purity verified by SDS-PAGE and Coomassie gel staining as well as Western blot (Supplementary Figure S3). Both antigens were determined for optimal plate coating concentrations: A) 2ug/ml prefgB; B) 3ug/ml gB(AD1). Net OD (absorbance 450nm) was attained by subtracting the OD of Ag- from the OD of Ag+. ELISA reactivity was considered positive if the net OD was greater than or equal to 0.2 as determined by GPCMV negative serum. Assays on pooled sera were conducted minimum of 3 times and mean titer determined. In addition, varying coating concentration of control carrier recombinant GST protein (Genscript) was tested for background using both GPCMV seropositive and seronegative control sera (Supplementary Figure S4).

#### 2.5.2 Western blots

Western blots were carried out on cell lysates separated by 10% SDS-PAGE under denaturing conditions and performed as previously described (17, 33). For western blots, anti-epitope tag primary antibodies were used at 1/1000: mouse anti-FLAG M2 (Sigma) (17, 19); mouse anti-GST (Genscript). Secondary antibodies: anti-mouse HRP-linked secondary antibodies for western blot (Cell Signaling) were used at 1/2000 (19). FLAG epitope was used to detect prefgB protein expression in transduced GPL cell monolayers. GST was used to detect gB(AD1)-GST tagged fusion protein. PrefgB was evaluated under denaturing and non-denaturing conditions. For evaluation under non-denaturing conditions prefgB was analyzed as previously described (20) with gB protein expression (monomeric) and multimerized protein both detected (Supplementary Figure S2).

### 2.6 Guinea pig interferon gamma enzyme-linked immunospot (ELISPOT) assay

The guinea pig IFNγ ELISPOT assays were performed following a previously described protocol using freshly isolated splenocytes and GPCMV GP123 peptide pools (19). Anti-guinea pig IFNγ monoclonal antibodies (V-E4 and N-G3) were previously characterized (54). Briefly, PVDF membrane 96 well plates were coated with guinea pig IFNγ capture antibody, V-E4, and incubated overnight at 4°C, blocked then freshly isolated splenocytes added before being exposed to GPCMV GP123 15mer peptide pools and incubated for 18 hrs. Biotinylated detection antibody, N-G3 was added before streptavidin-AP conjugate (R&D Systems) then detected with BCIP/NBT (Life Technologies). Membranes were dried before spots were counted on ImmunoSpot S6 (CTL). Final counts were calculated based on spot forming cells (SFC) per 10^6^ cells after background spots (cells only without any stimulation) were subtracted. Con A (10µg/ml) was used as an assay positive control and other controls included cells only control, DMSO control (peptide background), and nonspecific peptide control and media only control. Control animals included positive control (posC) GPCMV infected (n=3) and negative control (negC) uninfected animals (n=2) prior to peptide stimulation.

GP123 peptide pools. Eighty-seven 15mer peptides (overlapping by 11 amino acids) covering the unique IE1 protein coding sequence GP123 were generated by GenScript. Peptide pools included 8- 10 peptides generating a total of 19 pools. Each pool was tested as described above (GP IFNγ ELISPOT assays) to determine the most reactive pools.

### 2.7 Statistical analysis

All statistical analyses were conducted with GraphPad Prism (version 7) software. Fisher’s exact test, Student’s t test (unpaired) with significance taken as a p value of <0.05 or as specified in the figure legends.

## 3 Results

### 3.1 GPCMV convalescent immunity (22122) does not prevent reinfection (TAMYC strain)

In order to demonstrate the limited ability of natural GPCMV infection (22122 strain) to provide cross strain protection against horizontal (SQ) challenge (TAMYC strain), animals were infected with 22122 strain GPCMV to establish convalescent natural immunity. Seronegative animals (n=12) were inoculated with a single injection (10^5^ pfu, SQ) of GPCMV (22122 strain) to mimic natural infection. Animals were evaluated for anti-GPCMV seropositive status as well as evaluated for immune response to specific viral glycoprotein complexes (gB, gM/gN, gH/gL and PC) at 3 months post-infection. All animals seroconverted to GPCMV seropositive status (mean anti-GPCMV titer 5120). Animals also had an immune response to all GPCMV viral glycoprotein complexes (Supplementary Figure S1). Next, animals were challenged SQ with10^5^ pfu of GPCMV (TAMYC). At 4, 8, 12 and 27 dpi, animals were randomly euthanized (n=3) and viral load in target tissues evaluated (lung, liver, spleen and blood). Additionally, at 27 dpi, salivary gland tissue was also evaluated for viral load. Results (Figure 1) were compared to a previous historical study of TAMYC strain virus challenge with the same virus stock in seronegative animals infected with identical dose of virus (19) and demonstrated that TAMYC strain retained the ability to disseminate to all target organs, including salivary glands despite GPCMV (22122) seropositive status of animals. However, viral load at all time points was significantly reduced compared to previous TAMYC virus dissemination in seronegative animals except in the D27 spleen which was not significant (Figure 1) (25). Overall, results demonstrated the limited ability of 22122 strain convalescent immunity to prevent viremia and dissemination of heterologous TAMYC strain virus challenge.

**Figure 1.**
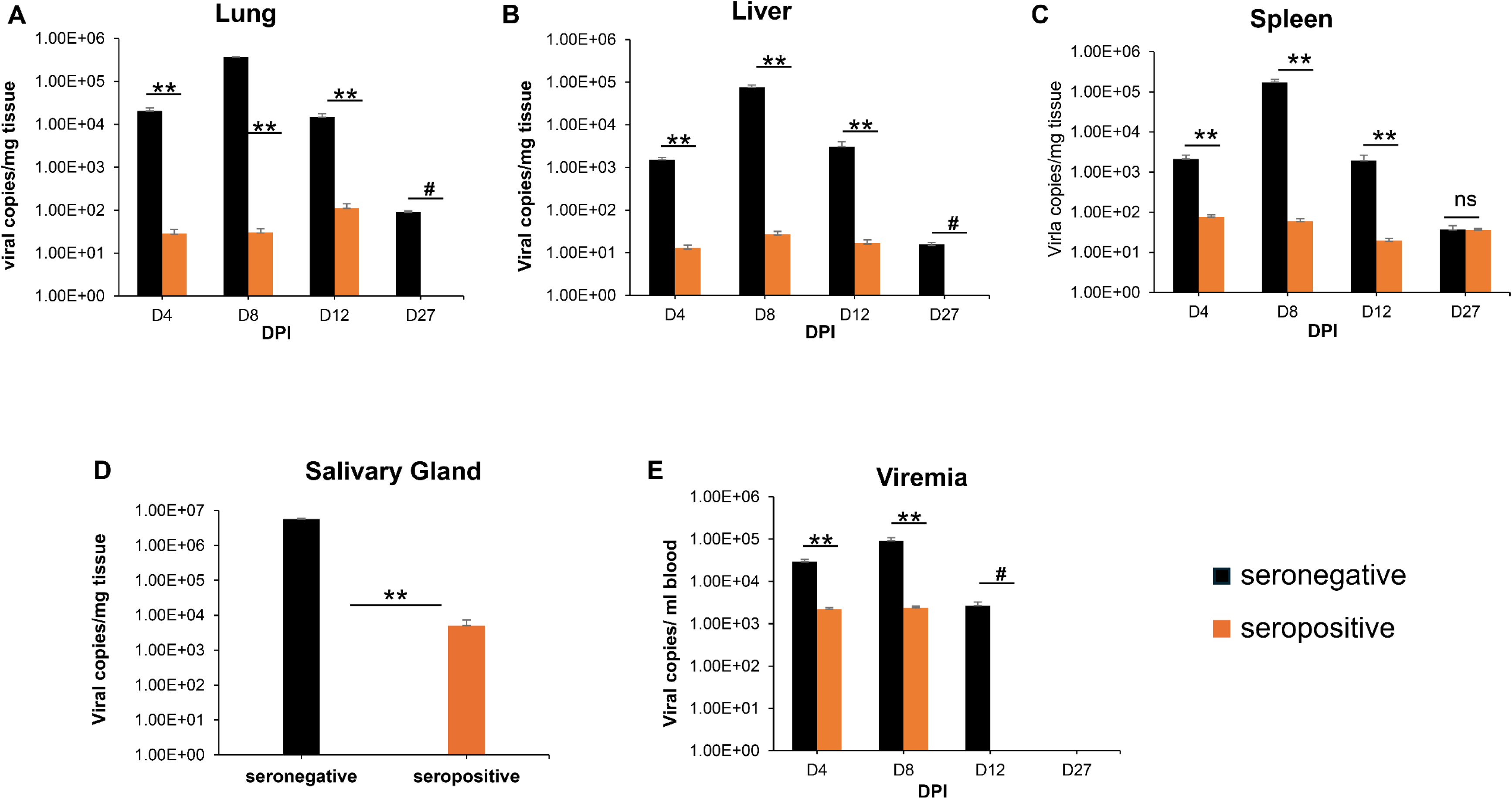
GPCMV (22122 strain) convalescent immunity does not protect against heterologous GPCMV (TAMYC strain) challenge. Seronegative animals (n=12) were inoculated once with 22122 strain GPCMV (10^5^ pfu, SQ). After verification of seroconversion, animals were designated as seropositive (orange). At 3 months post original infection, animals were challenged with TAMYC strain GPCMV (10^5^ pfu, SQ). A control group (n = 12) of seronegative animals (black) were similarly challenged with TAMYC strain GPCMV. At 4, 8, 12, and 27 days post infection (DPI), 3 animals per group were evaluated for viral load in target organs by real-time PCR of tissue extracted DNA. Viral load is plotted as the mean number of viral genome copies/mg tissue. Salivary gland tissue was evaluated only at day 27. (A) Lung; (B) liver; (C) spleen; (D) salivary gland. Viremia (E) at 4, 8, 12, and 27 dpi is plotted as the mean number of genome copies/ml blood. Statistical analysis determined by unpaired Student’s t test; ** *p* <0.001; ns = not significant; ^#^ viral load in CMV^+^ group below level of detection.

### 3.2 Preconception Ad vaccine (AdgB or AdgB+AdIE1) immune response in animals

Subsequently, the ability of Ad based vaccines to trimeric capable gB and IE1 antigens were evaluated for ability to provide cross strain protection against cCMV. An outline of the study is shown in Figure 2. Seronegative dams were randomly assigned to two different vaccine groups: Group 1 (AdgB), n=17; Group 2 (AdgB+ AdIE1), n=13. At days 0, 28 and 56, animals were vaccinated SQ with a specific CMV vaccine candidate according to assigned group (vaccine dosage 10^8^ TDU/shot per Ad vector) with bleeds at days 26, 54 and 78 for evaluation of anti-GPCMV antibody response. At day 78 post initial vaccination, antibody immune response was characterized in depth with a series of ELISA assays for individual animals: anti-GPCMV; anti-gB(wt); prefusion gB; and gB (AD-1 domain). Results were compared between groups 1 and 2 and also with pooled convalescent sera from single shot 22122 infected animals (Figure 3). The pooled sera from single shot GPCMV infected animals had similar anti-GPCMV mean titer to vaccine groups 1 and 2 but the anti-GPCMV titer for individual animals in the AdgB vaccine group were more varied compared to the AdgB+AdIE1 group despite similar overall mean titer (Figure 3A). The anti-gB response (trimeric capable gB) was on average higher for group 1 animals (mean titer 12047) than the AdgB+AdIE1 group (mean titer 4923) which was statistically significant (Figure 3B), but these titers were not uniformly higher for all animals in group 1 compared to group 2. Both groups had statistically higher anti-gB titer compared to pooled sera from single shot 22122 strain infected animals.

**Figure 2.**
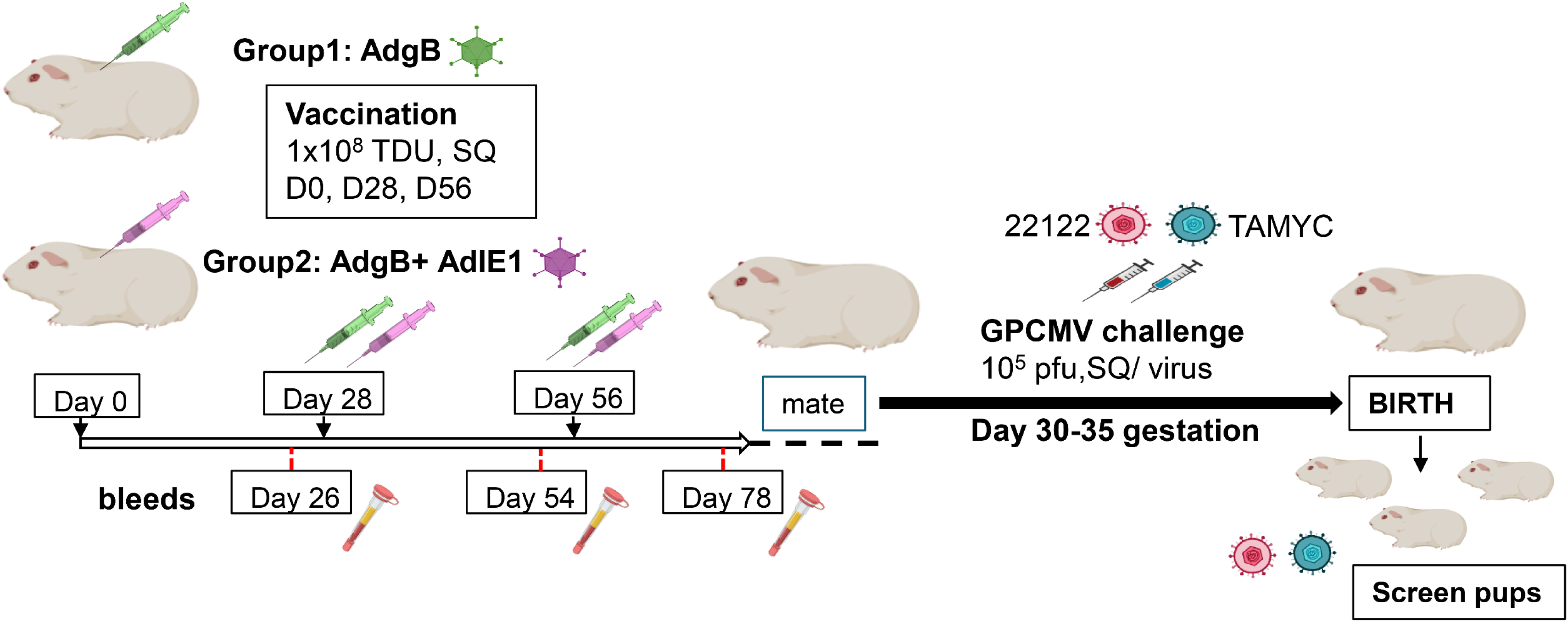
Schematic overview of preconception vaccine study and GPCMV congenital virus challenge with dual viral strains (22122 and TAMYC). Seronegative dams: Group 1 (AdgB, n=17) or Group 2 (AdgB+AdIE1, n=13) were vaccinated 3 times (10^8^ TDU) via SQ route on days 0, 28, and 56. Serum was collected 3 to 4 weeks after each vaccination (days 26, 54, and 78). The dams were mated, and during the late second trimester of pregnancy, the animals were challenged with both 22122 and TAMYC strains (10^5^ pfu/virus, SQ route) then followed to term. Viral load in the target organs (liver, lung, spleen, brain) of live-born or still-born pups was evaluated by real-time PCR. Placenta tissue when available was also evaluated for viral load. A control group of unvaccinated pregnant dams (n=10) were similarly challenged with GPCMV and pup tissue evaluated for viral load.

**Figure 3.**
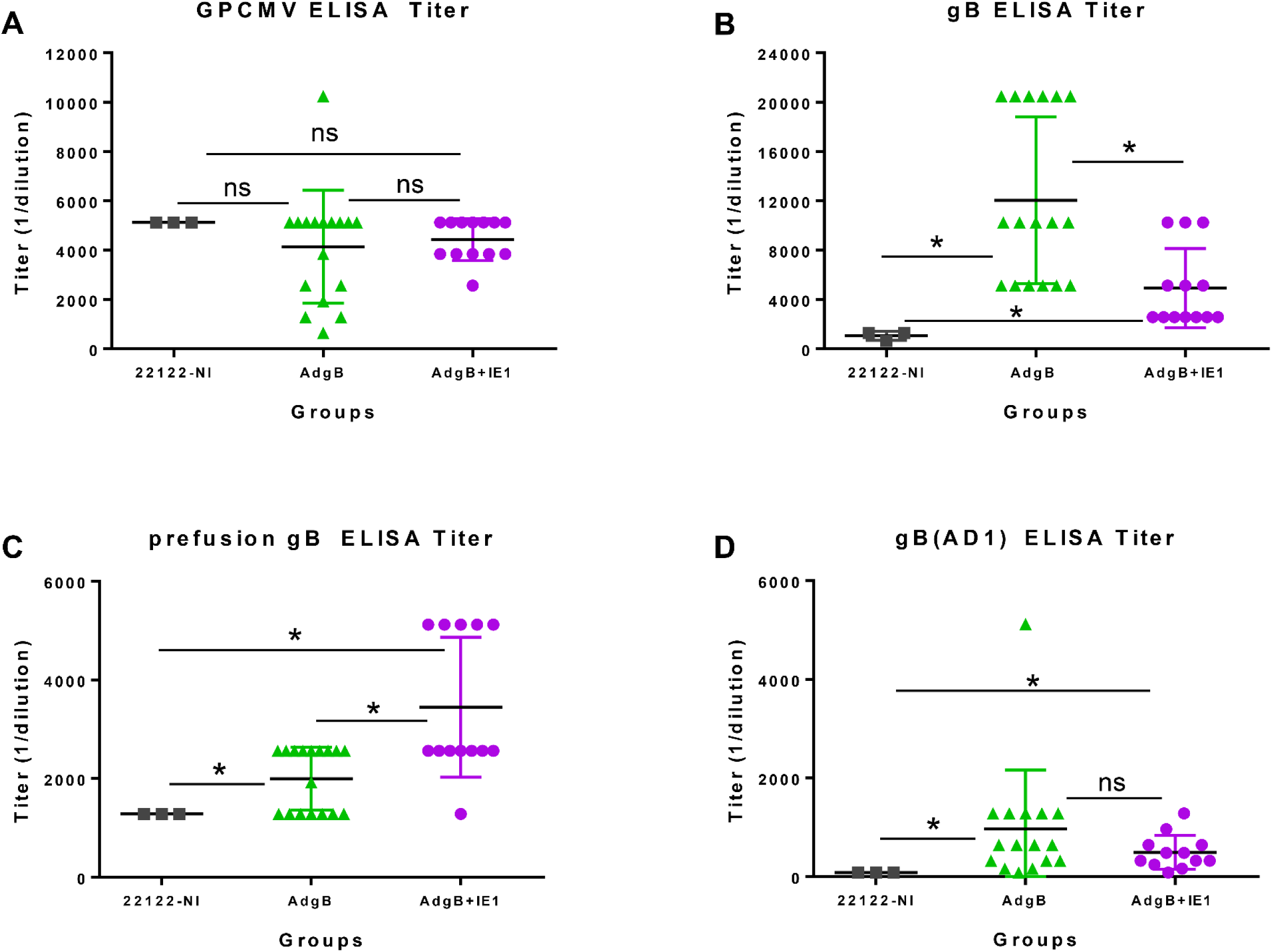
Antibody immune responses to AdgB or AdgB+AdIE1 vaccines compared to convalescent immunity of wildtype GPCMV naturally infected (22122-NI) animals. Individual animal sera of AdgB (green) or AdgB+AdIE1 (purple) vaccinated animals were compared to pooled convalescent sera of naturally infected GPCMV (black) animals for antibody titers to specific target ELISAs: (A) anti-GPCMV; (B) anti-gB; (C) anti-prefusion gB; or (D) anti-gB(AD1). Pooled 22122- NI sera assays were repeated 3 times. Mean value from each group represented by black horizontal line. Statistical analysis was determined by unpaired Student’s t test, \**p* < 0.05; ns= not significant.

Prefusion gB is a newly defined antigen for HCMV (52) and not previously evaluated for GPCMV. Based on BLAST alignment between HCMV gB and GPCMV gB predicted amino acid sequence, a locked version of GPCMV prefusion gB was generated by incorporation of specific codon changes used for generation of HCMV prefusion gB and removal of the furin cleavage site (52). A synthetic codon optimized gene encoding prefusion gB (prefgB) was generated (Genscript) and cloned into a mammalian expression plasmid used to generate ELISA coating prefgB antigen as described in materials and methods. Prefusion gB expression in plamid transfected cells was verified by western blot assay which detected both monomeric and multimeric forms of prefgB (Supplementary Figure S2). Evaluation of the immune response to prefusion gB indicated that group 2 animals had a higher titer (mean titer 3446) compared to group 1 animals (mean titer 1995) and was significantly different between the groups (Figure 3C). Both vaccine groups had significantly higher anti-prefusion gB titers compared to pooled sera from 22122 infected animals (mean titer 1280), Figure 3C.

The gB AD1 region is considered to be the immunodominant domain in HCMV gB but not previously evaluated for GPCMV in prior GPCMV gB vaccine studies. A bacterial expression plasmid encoding GST-gB(AD1) fusion protein was based on BLAST alignment of HCMV gB AD1 domain with GPCMV gB ORF (Supplementary Figure S3A) (53). A synthetic gene encoding an extended GPCMV gBAD1 fused to the C-terminus of GST carrier protein was generated as described in materials and methods. GST-gB(AD1) protein was expressed in bacteria and recombinant protein purified by GST affinity column chromatography and verified by Coomassie stained SDS-PAGE (Supplementary Figure S3B) and Western blot analysis (Supplementary Figure S3C) prior to use as gB(AD1) ELISA coating antigen. A control ELISA with recombinant GST protein (Genscript) lacked activity with GPCMV seropositive and seronegative sera demonstrating a lack of any preexisting antibodies to GST protein in animals (Supplementary Figure S4). Anti-gB(AD1) ELISA immune responses were more tightly clustered for individual animals within vaccine groups with only one animal titer relatively high in group 1 (5120) compared to mean value of 965. However, the difference in mean titer between group 1 and group 2 animals was not significant (Figure 3D). Both vaccine groups produced a significantly higher anti-gB(AD-1) response compared to 22122 strain convalescent sera (mean titer 80), Figure 3D. Overall, based on ELISA studies, the biggest difference between vaccine groups was in the immune response to prefusion gB, which was highest in group 2 animals and also for gB but the response was highest in group 1 animals by greater than 2-fold.

Next, sera from vaccinated animal groups or convalescent sera (22122 infected animals) were evaluated for neutralizing antibody titers on both fibroblast (GPL) and epithelial (REPI) cells for GPCMV strains 22122 and TAMYC (Figure 4). Evaluation of GPL NA_50_ (22122 strain) for individual sera from each vaccine group compared to pooled convalescent sera did not demonstrate any significant difference between groups but individual NA_50_ titers for group 1 animals was more widely spread compared to group 2 animals (Figure 4A). Pooled sera within groups were subsequently used for evaluation of NA_50_ (22122) on epithelial cells where group 1 vaccine had a significantly higher NA_50_ titer compared to group 2 or 22122 infected animal sera (Figure 4B). There was no significant difference in NA_50_ titer between group 2 animals and 22122 convalescent sera.

**Figure 4.**
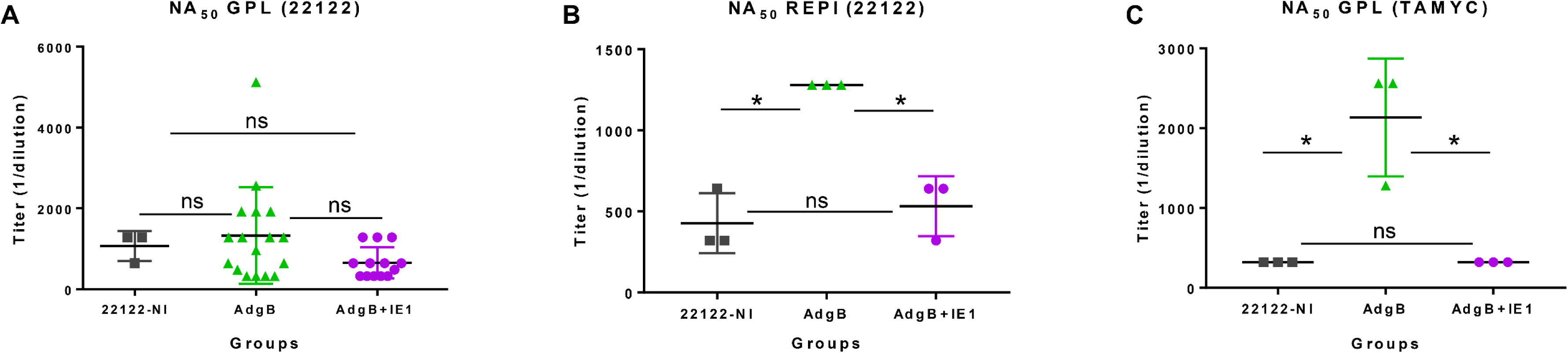
GPCMV neutralization by anti-AdgB or anti-AdgB+AdIE1 animal sera compared to GPCMV. (**22122**) **natural convalescent sera (22122-NI).** Individual animal sera of AdgB (green) or AdgB+AdIE1 (purple) vaccinated animals were compared to pooled sera of GPCMV (22122) convalescent sera (black) animals for neutralization of 22122 or TAMYC strain GPCMV on different cell lines: (A) neutralization of 22122 on GPL fibroblasts cells; (B) neutralization of 22122 on epithelial cells (REPI); (C) neutralization of TAMYC virus on GPL cells. Pooled 22122-NI sera assays were repeated 3 times. Mean value from each group represented by black horizontal line. Statistical analysis was determined by unpaired Student’s t test, \**p* < 0.05; ns= not significant. Assays were carried out with pooled sera within different groups except for (A) where vaccine sera from individual animals from both vaccine groups was evaluated.

Evaluation of NA_50_ (TAMYC) resulted in higher NA_50_ titer for the AdgB (group 1) vaccine group compared to other sera groups (Figure 4C). Overall, results suggest that group 1 (gB vaccine) sera had slightly better neutralizing capability against both strains of virus.

Group 2 vaccinated animals (AdgB+AdIE1) were additionally evaluated for cell mediated response against IE1 antigen utilizing a guinea pig IFN-γ ELISPOT assay previously developed (48). An overlapping peptide library to GP123 (IE1 unique exon coding region) was used to evaluate splenocytes isolated from vaccinated animals (n = 3) for induction of a cell mediated response to identify three different active peptide pools containing eight to ten 15mer peptides corresponding to GP123. Vaccinated animals demonstrated an immunogenic response to IE1 (Figure 5). Cellular response in vaccinated animals was similar to GPCMV infected positive control (posC) animals (n = 3). Negative control (negC) uninfected animals showed no response to GP123 peptide stimulation.

**Figure 5.**
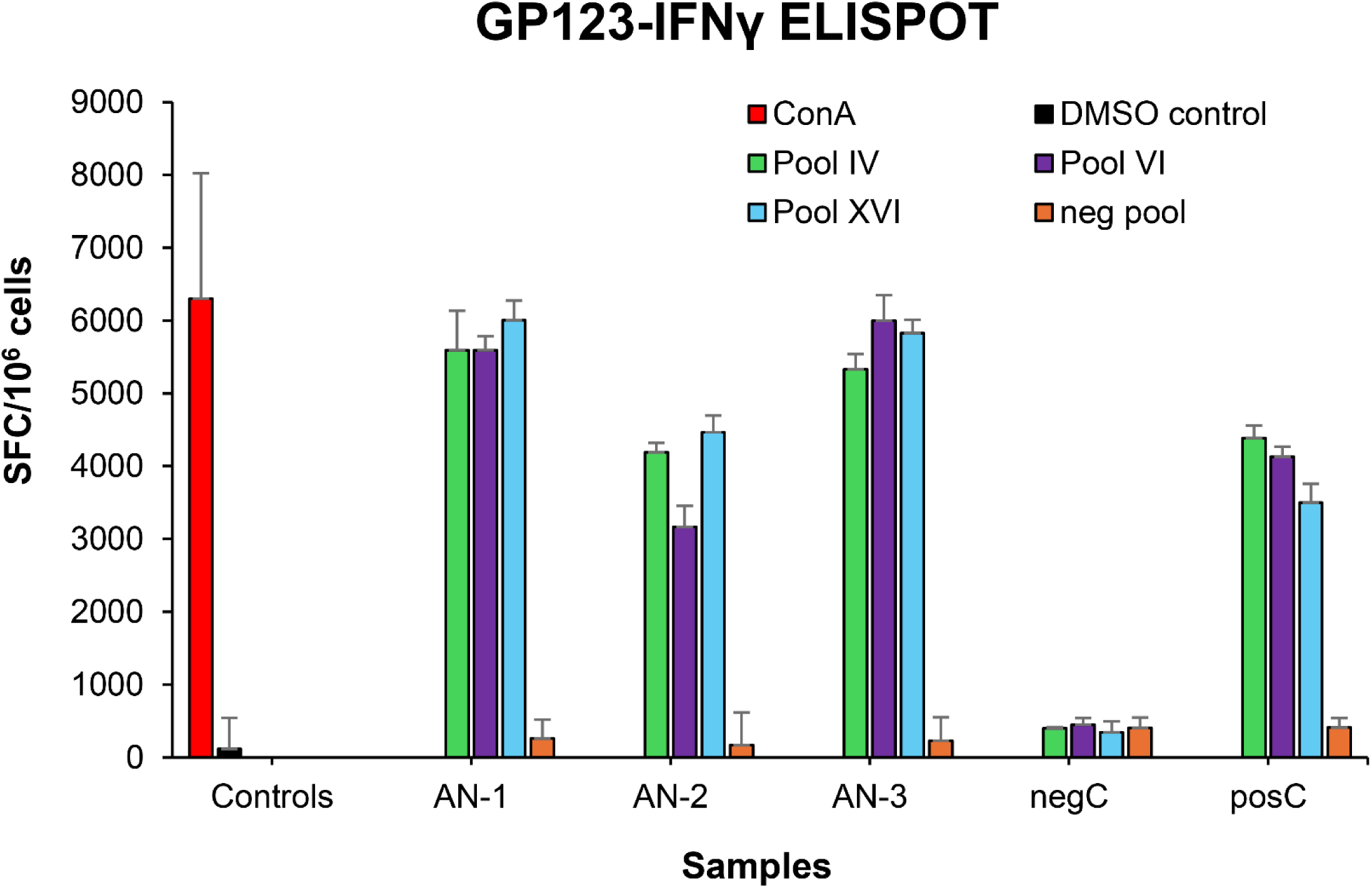
T cell response to GPCMV IE1. Guinea pig specific IFNγ ELISPOT was performed using pools overlapping peptides to GP123. Three GP123-reactive peptide pools, IV, VI, and XVI (green, purple and blue bars) reacted with primed splenocytes from AdgB+AdIE1 vaccinated animals (AN-1, 2,3), GPCMV primed (posC, n=3) or uninfected controls (negC, n=2). ConA assay control stimulation, red; DMSO control, black; unresponsive GP123 negative peptide pool, orange. Final counts were calculated based on the number of spot-forming cells (SFC) per 10^6^ cells after background subtraction.

Results demonstrated that co-vaccination with AdgB+AdIE1 did not impair induction of IE1 cell mediated response in group 2 animals.

### 3.3 Preconception vaccine protection against cCMV

At approximately 4 weeks post last vaccination, dams were mated with seronegative males. At 30–35 days of pregnancy (late second 2nd trimester), dams were challenged with wild type GPCMV. At the time of challenge, the average duration since last vaccination was approximately 3 months. Challenge virus consisted of two strains of GPCMV (22122 and TAMYC strains) with each strain being separately administered into opposite flanks of the animal by SQ injection (10^5^ pfu GPCMV per injection). A control group (group 3) of seronegative unvaccinated dams (n = 10) were mated and similarly challenged with wild type GPCMV strains during 2nd trimester. Animals were allowed to proceed to term, and pups from all three groups were evaluated for viral load. Results are shown in Tables 1 & 2 and Figure 6. A summary table of the congenital CMV study outcome is shown in Table 1, which compares litters from vaccine groups 1 and 2 to those from unvaccinated control group 3 pups. In the control unvaccinated group, 25% (9/36) of animals were stillborn compared to one stillborn pup in each vaccine group representing 2% (group 1) and 2.8% (group 2) of total litter. In the unvaccinated group, 78% of stillborn pups in the control group were CMV positive but none of the stillborn pups in either vaccine group had detectable virus in tissue organs. Despite the absence of detectable virus in stillborn pup tissues in the vaccine groups, we assumed that it is probable that maternal CMV infection was the basis for the pup death and not a complication of pregnancy.

**Figure 6.**
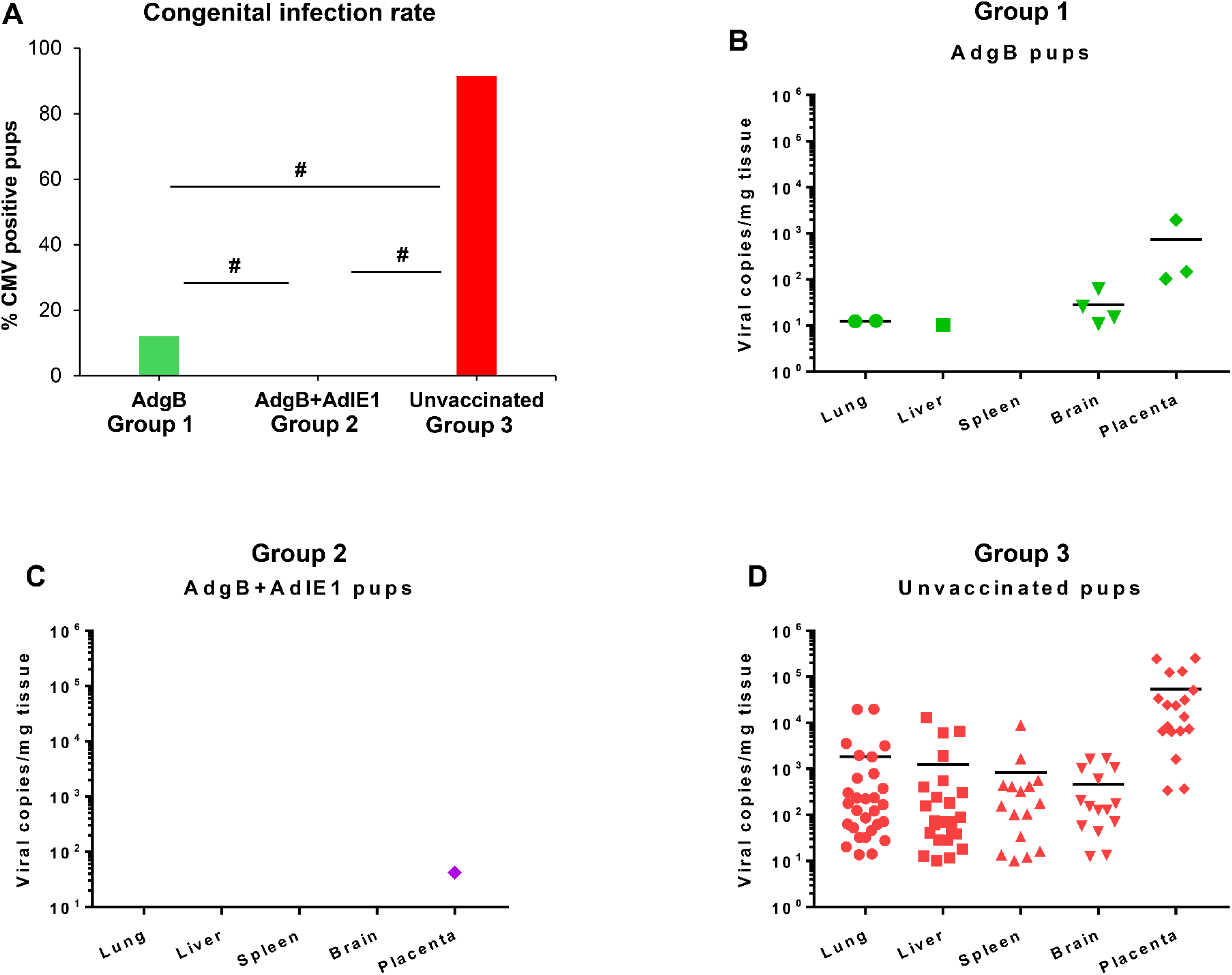
Vaccine protection against dual strain GPCMV congenital infection. (A) Congenital infection rate determined by percent GPCMV positive pups in AdgB vaccinated (12%, green) or AdgB+AdIE1 vaccinated (0%, purple) compared to unvaccinated (92%, red) pups based on detectable virus in tissues tested by real-time PCR. Statistical analysis performed by Student t-test # *p*< 0.05. Pup viral load in each study groups (B) AdgB vaccinated (Group 1, green); (C) AdgB+AdIE1 vaccinated (Group2, purple); or (D) Unvaccinated group (Group 3, red) with detectable levels of GPCMV found in lung, liver, spleen, brain or placental tissues. Total number of detectable samples from each group and statistical analysis represented in Table 2. Mean values from each test group represented by black horizontal line.

**Table 1.** Congenital infection outcomes for live versus still-born pups.

|  | **Group 1:**<br>**AdgB** | **Group 2:**<br>**AdgB+AdIE1** | **Group 3:**<br>**Unvaccinated** |
| --- | --- | --- | --- |
| Pregnant | 17/17 (100%) | 13/13 (100%) | 10/10 (100%) |
| Litters delivered | 17 | 13 | 10 |
| Litters with only live pups | 16 | 12 | 7 |
| Litters with mix (live & dead) pups | 1 | 1 | 1 |
| Litters with only dead pups | 0 | 0 | 2 |
| Total pups (live) | 49 (98.0%) | 35 (97.2%) | 27 (75%) |
| *Live pups CMV+* | *6/49\*# (12.2%)* | *0/35\*# (0.0%)* | *26/27\* (96.3%)* |
| Total pups (dead) | 1 (2.0%) | 1 (2.8%) | 9 (25%) |
| *Stillbirth pups CMV+* | *0/1 (0.0%)* | *0/1 (0.0%)* | *7/9 (77.8%)* |

\*Statistical analysis comparing GPCMV positive pups of live-born in AdgB or AdgB+AdIE1 vaccinated against unvaccinated groups: \**p* <0.0001 determined by Fisher exact test #Comparison of GPCMV positive pups of live-born between vaccine groups AdgB vs Adgb+AdIE1groups: #*p* < 0.05 determined by Fisher exact test
\*Statistical analysis comparing GPCMV positive pups of live-born in AdgB or AdgB+AdIE1 vaccinated against unvaccinated groups: \**p* <0.0001 determined by Fisher exact test

**Table 2.** cCMV outcome of pups with detectable virus in organs

|  | **Group 1:**<br>**AdgB\*** | **Group 2:**<br>**AdgB+AdIE1\*** | **Group 3:**<br>**Unvaccinated\*** |
| --- | --- | --- | --- |
| **CMV+ pups** | **6/50<sup>#</sup> (12.0%)** | **0/36<sup>#</sup> (0%)** | **33/36 (91.67%)** |
| Lung | 2/50 (4.0%) | 0/36 (0.0%) | 29/36 (80.56%) |
| Liver | 1/50 (2%) | 0/36 (0.0%) | 24/36 (66.67%) |
| Spleen | 0/50 (0%) | 0/36 (0.0%) | 15/36 (41.6%) |
| Brain | 4/50<sup>□</sup> (8.0%) | 0/36<sup>□</sup> (0.0%) | 15/36 (41.6%) |
| Placenta | 3/20 (15%) | 1/25 (4.0%) | 18/18 (100%) |

\*Significant difference between each study group (AdgB or AdgB+AdIE1) compared to unvaccinated group in total CMV+ pups and each tissue groups. Statistics determined by Fisher’s exact test $p < 0.001$
<sup>#</sup>Significant difference in CMV+ pups between AdgB and AdgB+AdIE1 groups determined by Fishers exact test $p < 0.05$ )
<sup>□</sup>No significant difference in all tissues except in the brain between the two study groups (1 & 2) as determined by z test $p < 0.05$

Additionally, we cannot rule out the possibility of virus being present in pups at below detectable assay levels. Both vaccine groups had approximately similar high percentages of live pups (98% and 97.2% for groups 1 and 2 respectively) compared to 75% live pups in the no vaccine control group. A comparison in the numbers of CMV positive live pups was statistically significant between vaccine groups and no vaccine control (Table 1): control vaccine group 3 live pups (96.3% CMV+ pups); AdgB vaccine group 1 (12.2% CMV+ pups); the AdgB+AdIE1 vaccine group 3 (0% CMV+ pups).

Furthermore, there was a statistically significant difference in CMV positive live pups between the two study groups (group 1 vs group 2).

Overall, both vaccine strategies were highly successful in reducing cCMV transmission rates to pups (Table 2). However, the combined gB+IE1 vaccine strategy was most successful with 0% detectable virus in all pups compared to 91.7% of pups positive in the control group 3, see Table 2 and Figure 6. The gB vaccine strategy was also highly effective with reduction in cCMV transmission to 12% in pups in group 1, Table 2 and Figure 6. Both vaccine groups had statistically significant outcomes in reduction of cCMV transmission. In the no vaccine control group 3 pups, virus disseminated to all target organs tested (Table 2 and Figure 6), with 41.6% pups positive for CMV in the brain compared to 8% in the gB vaccine group and 0% in the gB+IE1 vaccine group (Table 2). In both vaccine groups, virus was detected in the placenta at a low level, 1/25 pups (4%) for group 2 and 3/20 (15%) for group 1, but this was a substantial decrease from the level of virus in the placentas of the no vaccine control group,18/18 (100%), see Table 2 and Figure 6.

## 4 Discussion

Previous GPCMV vaccine studies have demonstrated the limited ability of gB to provide high level protection against CMV, especially for cross strain protection against highly cell associated clinical GPCMV strain (TAMYC). Potentially, inclusion of a T cell antigen can enhance cross strain protection against cCMV. In HCMV, convalescent patients produce T cell responses against two specific viral proteins (pp65 and IE1) of significance for vaccine development (45) and GPCMV infected animals exhibit T cell response to both IE1 and pp65 homologs (21, 34, 48). Multiple studies with GPCMV pp65 (GP83) or in combination with gB did not enhance cCMV protection against 22122 strain, reviewed in Choi & McGregor 2025 (12). Indeed complete protection against cCMV (22122 strain) can be achieved by use of a live attenuated GPCMV strain lacking pp65 antigen (24). Our previous IE1 vaccine study (48) demonstrated the ability of cell mediated T cell response to provide high level cross strain protection against cCMV. This present study demonstrated an ability of gB immune response to synergize with IE1 response in the gB+IE1 vaccine group to further improve cCMV protection unlike previous gB and pp65 dual antigen vaccine strategies. IE1 cell mediated response was unaffected by the inclusion of gB antigen. Importantly, the gB+IE1 vaccine was cross strain protective against cCMV with complete protection. In contrast, a gB vaccine reduced cCMV transmission to 12% compared to 92% in the no vaccine control group (80% reduction). However, the AdgB vaccine lacks cross strain protection against TAMYC strain challenge in a non-pregnant animal model with the ability to fully disseminate in vaccinated animals (23). An earlier IE1 vaccine strategy reduced cCMV transmission to 23% (48). A previous Ad vaccine based GPCMV (22122) study by Inoue and colleagues (55) with a recombinant Ad gB (monomeric) vaccine strategy resulted in a 13% cCMV transmission rate to pups in the gB vaccine group compared to 75% cCMV transmission in a control AdlacZ vaccine group, a reduction in transmission of 62% against GPCMV (22122) indicating that a trimeric gB further enhances protection against cCMV compared to monomeric gB. Importantly in the Inoue study (55), the cCMV transmission rate in the control vaccine group (AdlacZ) was generally similar to that of unvaccinated animals in other 22122 strain cCMV vaccine studies (12). Additionally, Ad based gB and pp65 vaccine strategies do not protect against TAMYC strain challenge in a non-pregnant model (21, 23). This indicates that the Ad vector platform does not contribute to the protection against GPCMV but is represents a simple but effective strategy for target antigen expression. Indeed, an Ad based vaccine has demonstrated efficacy against congenital Zika virus in preclinical studies (56) and Ad based vaccines have been used successfully in the clinic against Ebola virus and SARS-CoV2 (57, 58). In future studies, an oral vaccination route is a potentially safer approach (59) and avoids pre-existing immunity to specific Ad virus serotypes as well as low level risk of blood clots from i/m vaccination (60, 61).

Guinea pig T cell response to IE1 was evaluated with a guinea pig specific IFNγ ELISPOT assay using 15mer overlapping peptides covering the entire length of GP123 to identify three reactive pools of peptides. Each of the 15mer peptides overlapped on GP123 protein by 11 amino acid to comprehensively cover potential antigenic peptides and peptides were used at a concentration of 5µg/ml which has been shown to be just as effective as using 9mers at a lower concentration (62). A limitation of available immunological reagents dedicated to the guinea pig generally prevents the ability to further characterize guinea pig cell mediated response much beyond IFNγ ELISPOT assays and additional dedicated guinea pig T cell assays need to be developed. Studies in transplant patients suggest the importance of CD8^+^ T cell response against IE1 for protection from CMV (63) and our assumption is that AdIE1 vaccine elicits a CD8^+^ T cell response. HCMV (Towne) vaccination of seronegative patients resulted in IE1 CD8^+^ T cell response but not pp65 (64). A HCMV Toledo/Towne vaccine similarly induced IE focused response (65). Additionally, CD8^+^ T cell IE1 response but not pp65 is associated with protection against HCMV in solid organ transplant patients (63). HCMV studies in human placental decidual tissue and resident T cells suggest the importance of HCMV specific resident CD8^+^ T cells for protection against HCMV (66). Evaluation of resident guinea pig placental T cells would likely be informative especially if vaccine responses were contrasted with natural convalescent immunity. Future GPCMV research ideally should better define CD4^+^ and CD8^+^ T cell responses to GPCMV IE1 as well as other viral antigens. Potentially, a transcriptomics approach applied to HCMV clinical studies that enables characterization of signature T cell response to a HCMV vaccine could be applied to GPCMV vaccines (67). The recent in-depth sequencing of the Hartley strain guinea pig genome enables an RNA seq transcriptomics approach in the guinea pig (68). This strategy has recently been successfully applied to CD4^+^ and CD8^+^ T cell studies and HSV vaccine research in this animal model (69).

A limitation of previous gB vaccine studies in the guinea pig model is that the majority of gB antigens studied lacked the ability to form a trimeric gB complex, which enhances the level of neutralizing antibody response against GPCMV (20). Although a trimeric gB vaccine was highly effective against 22122 strain in a horizontal (SQ) challenge model, it lacked cross strain protection against GPCMV (TAMYC) dissemination (SQ) despite 99% similarity in gB amino acid sequence between strains (20, 21). This demonstrated the limitation of gB cross strain protection despite success against cCMV with approximately 50% protection against cCMV (22122) in subunit and vector based gB vaccine strategies, reviewed in Choi and McGregor 2025 (12). The immunological basis for the limitation of prior GPCMV gB vaccine strategies in the guinea pig model are poorly defined. In this report, additional aspects of the gB immune response were characterized between vaccine groups and in comparison to natural convalescent GPCMV immunity (22122 strain) by using novel gB specific ELISAs which included prefusion gB and gB(AD1) as well as established gB and anti-GPCMV ELISAs (20). In HCMV, the gB AD1 domain is necessary for oligomerization of gB (53) and presumably the case for GPCMV. HCMV gB AD1 domain antibody response is common in all convalescent patients and AD1 is also a major target for neutralizing antibodies (70–72). It is interesting to note that in HCMV clinical trials for both gB/MF59 and Moderna mRNA-1647 vaccines, gB AD1 binding antibodies were not detected (73, 74). In this current GPCMV vaccine study all vaccinated animals had a specific response to gB(AD1) antigen, as did convalescent sera (22122 strain), see Figure 1. Both vaccine groups induced a higher titer anti-gB response compared to natural immunity with the gB vaccine group inducing approximately 2-fold higher titer response compared to gB+IE1 vaccine group but individual responses were more wide spread (Figure 3). In contrast, the AD1 immune response in both vaccine groups was more tightly clustered where both groups had statistically higher titers than natural convalescent immunity.

HCMV gB is the fusogenic protein and has the potential to exist in a prefusion confirmation on viral particles which might be more effective as a vaccine target and a locked version of HCMV prefusion gB was recently described (52, 75). Based on alignment of HCMV and GPCMV gB amino acid sequences, similar modifications were made to the GPCMV gB ORF to generate a synthetic gene encoding a locked prefusion gB. Both vaccine groups exhibited a response to the GPCMV prefusion gB antigen with antibody titers statistically higher than natural immunity but the gB+IE1 vaccine group had a higher response compared to the gB only vaccine. Results suggest that the presence of IE1 antigen was not impactful on gB immune response to AD1 domain despite lower anti-gB titer but that inclusion of IE1 antigen resulted in higher levels of anti-prefusion gB response. Potentially, this contributed to the enhanced protection against cCMV in the gB+IE1 vaccine group compared to gB group. In the case of one litter (dam#3) in the gB vaccine group, differential antibody immune response might have contributed to placental and pup infection. In this litter (dam#3), 2/3 pups and 2/3 placenta were infected with virus, the dam had a higher anti-gB titer (20480) and the highest gB(AD1) titer (5120) but a lower prefusion gB titer (1280) compared to the mean value, Figure 3. This suggests that neither a high AD1 nor a high gB antibody titer is not necessarily a predictor of positive outcome against virus challenge. In contrast, a low response to prefusion gB antigen might indicate a greater risk for cCMV. However, this aspect likely requires further evaluation of both AD1 and prefusion gB immune responses, but these studies are beyond the scope of this initial research.

Several antigenic domains exist for HCMV gB (76) and this is likely the case for GPCMV and future studies should also characterize potential additional homolog antigenic domains, especially a homolog N-terminal AD2 domain where only 50% of HCMV convalescent patients have an antibody response to this neutralizing antibody domain (76, 77).

In HCMV, antibodies to gB as well as non-neutralizing antibodies to other viral proteins can provide protection by ADCC and ADCP pathways (73, 78, 79) but currently evaluation of this aspect of the immune response is missing in guinea pig studies. Most certainly, GPCMV monomeric gB in comparison to trimeric gB produces high level of non-neutralizing antibodies (20) and control of infection by ADCC/ADCP pathways is perhaps a realistic possibility. However, in animals, a gB vaccine lacks cross strain protection against a highly cell associated clinical TAMYC strain suggesting a limitation of this antibody cell associated immune response in the guinea pig (23). This is also suggested by the fact that natural GPCMV (22122) convalescent immunity with both neutralizing and non-neutralizing antibodies to various viral antigens fails to prevent infection by heterologous TAMYC strain challenge (Figure 1). Although antibody based ADCC effects may enable targeting of infected cells, HCMV has the capacity to evade ADCC induction (78–80).

Consequently, the importance of ADCC/ADCP pathways in contributing to protection against GPCMV is ambiguous especially if the virus has the capacity to evade these pathways as suggested by the ability of clinical strain TAMYC to evade both gB and GPCMV DISC vaccines which are highly effective against 22122 strain (23, 24).

An additional important aspect of antibody response that remains to be evaluated relates to fetal protection by transplacental transfer of protective maternal antibodies to the fetus in utero (81). In guinea pigs, both human and guinea pig IgG can be transferred across the placenta (82). Previously, anti-gB passive antibody therapy in guinea pigs demonstrated partial placental and fetal protection against GPCMV, which suggests that this is a potential protective strategy but transplacental transfer of gB antibodies to the fetus remains to be more fully investigated (83). Future evaluation in this model of transplacental transfer of antibodies is further merited by recent research, which indicates that HCMV antibodies in humans can activate fetal NK cells via Fc receptor enabling cytotoxic targeting of virus infected cells (84). Consequently, it would be important to determine in future research if this mechanism exists in the guinea pig model to improve the translational impact of GPCMV vaccine studies. Potentially, GPCMV gB protein is a T cell target antigen based on studies with HCMV gB in convalescent patients (85) and the possible contribution of cell mediated T cell response to vaccine protection should be evaluated in future GPCMV gB research. However as previously demonstrated, the lack of cross strain protection of an AdgB vaccine against highly cell associated TAMYC virus with 99% amino acid gB identity between strains suggests a T cell response to gB has minimal protective impact compared to antibody response (23).

In summary, a combined Ad based CMV vaccine strategy of gB and IE1 provides complete cCMV protection against both prototype 22122 strain and a novel clinical strain of GPCMV (TAMYC). There is a risk of cCMV from both primary and non-primary infection by a new CMV strain. Consequently, a vaccine strategy needs to both provide high efficacy and cross strain protection. Complete cross strain protection against cCMV is an important milestone in this model. Results suggest that a combined approach of CMV antibody (gB) and T cell (IE1) antigen vaccine candidates is an important foundational strategy and would similarly be highly protective against HCMV. However, a HCMV vaccine would possibly require the modification of IE1 to attenuate functional activity to improve vaccine safety. Potentially, results suggests that an antibody response against the viral PC, important for endocytic infection of non-fibroblast cells, is not absolutely required for protection against cCMV. However, a limitation of the current study is that virus challenge was by SQ route and did not evaluate horizontal animal to animal transmission. Additional inclusion of PC antigen in vaccine design would increase mucosal protection by oral/nasal route of virus infection in future studies.

## 6 Author Contributions

K. Yeon Choi: Conceptualization, Methodology, Investigation, Data curation, Formal analysis, Writing - original draft, Writing - review & editing. Yushu Qin: Methodology, Investigation, Data curation, Formal analysis, Writing – original draft, Writing - review & editing. Nadia S. El-Hamdi: Conceptualization, Methodology, Investigation, Data curation, Formal analysis, Writing – original draft, Writing - review & editing. Alistair McGregor: Conceptualization, Methodology, Investigation, Data curation, Formal analysis, Writing - original draft, Writing - review & editing, Funding acquisition.

## Acknowledgments

The authors would like to thank Austin Selman and Alex Nguyen for their excellent technical assistance. Hybridoma cell lines for the anti-IFN-γ monoclonal antibodies were a generous gift from Dr. Schäfer (Robert Koch Institute, Germany).

## 8 Supplementary Material

Supplementary Figures 1-4 see attached.

