## Supplementary material for "Complete cross strain protection against congenital cytomegalovirus infection requires a vaccine encoding key antibody (gB) and T-cell (immediate early 1 protein) viral antigens": Suppl Figures 1,2,3,4

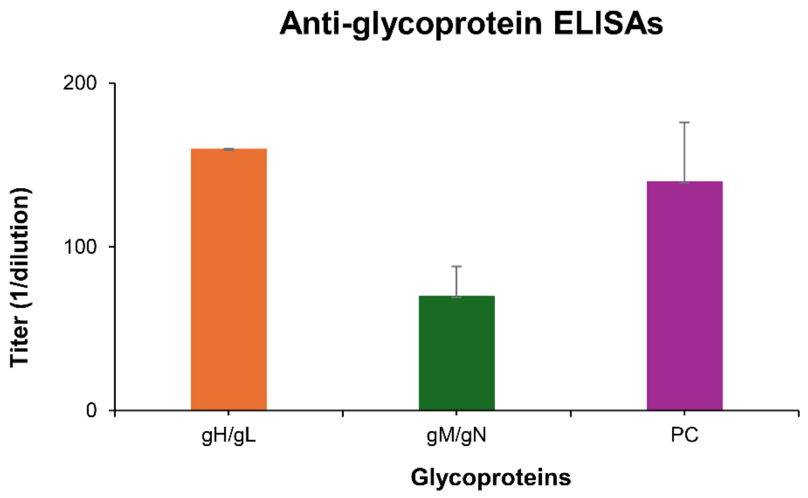


**Supplementary Figure 1. Immune response to single shot of 22122 strain GPCMV.** Specific viral glycoprotein complex ELISA assays were conducted on pooled sera from seropositive GPCMV(22122) infected animals and used to determine to determine ELISA titers for: anti-gHgL (orange); anti-gMgN (green); or anti-PC (purple) immune titers. Anti-GPCMV and anti-gB results are shown in Figure 3.

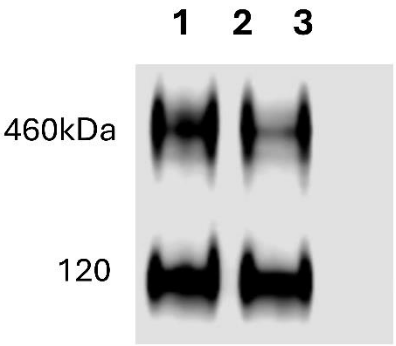


**Supplementary Figure 2. West blot analysis of GPCMV prefusion gB expression.** Prefusion gB transfected GPL cell lysate detected with anti-FLAG antibody (1/1000). Lane 1 & 2 prefgB plasmid GPL transfected cell monolayer total cell lysate at 24 hr post transfection; Lane 3 mock infected GPL cell lysate. SDS-PAGE carried out under non-denaturing conditions as described in materials and methods. Monomer prefgB (120 kDa) and triplex gB (460 kDa) indicated.

A

```
GPCMV  489  CLDQKRTAEVLHELSKINPSNILSAIFGVPVAARVVGDVISLAKCIEVNQSTVLIKGDMR  548
            C+DQ+RT EV  ELSKINPS ILSAI+  P+AAR +GDV+ LA C+ +NQ++V +   DM
HCMV   504  CVDQRRTLEVFKELSKINPSAILSAIYNKPIAARFMGDVLGLASCVTINQTSVKVLRDMN  563

GPCMV  549  KFSDDGKLEGCYSRPVVWFSMKNSTEVRLGQLGEDNEILLGTHRMETCQTQDYRIFVAGD  608
                G+    CYSRPVV F+   NS+ V+ GQLGEDNEILLG HR E CQ     +IF+AG+
HCMV   564  VKESPGR---CYSRPVVIFNFVNSSYVQYGQLGEDNEILLGNHRTEECQFPSLKIFIAGN  620

GPCMV  609  IGYEFQQYVFTKKINLSEIDIIDTMIALKTEPLENIDFKVLELYSRDELAQANVFDLESI  668
              YE+  Y+F + I+LS I   +D+MIAL  +PLEN DF+VLELYS+ EL  +NVFDLE I
HCMV   621  SAYEYVDYLFKRMIDLSSISTVDSMIALDIDPLENTDFRVLELYSQKELRSSNVFDLEEI  680
```

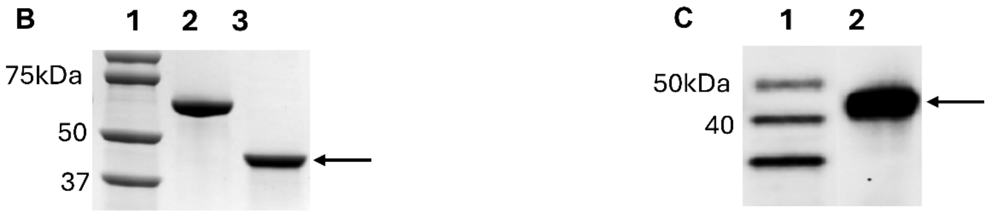


**Supplementary Figure 3. GPCMV gB(AD1) analysis.** (A) Protein BLAST alignment of HCMV gB AD1 compared to GPCMV gB AD1 region. Commercial HCMV (Towne) GSTgB(AD1) protein (Sigma), region 549-650 highlighted in blue. Aligned HCMV/GPCMV gB AD1 region as described by Britt et al., 2005 [53]. Underlined sequence GPCMV gB AD1 extended region used for generation of GPCMV GST-gB(AD1) fusion ORF with similar flanking sequence to commercial (Sigma) HCMV gB(AD1) fusion GST protein . (B) SDS PAGE gel of purified recombinant GPCMV GST-gB(AD1) protein (Genscript). Lane 1 marker; lane 2 BSA protein control; 3 GST-gB(AD1) protein resolved by Coomassie blue staining. (C) Western blot of recombinant GPCMV GST-gB(AD1) protein detected with anti-GST antibody. Lane 1 kDa size markers; lane 2 GST-gBAD1 protein.

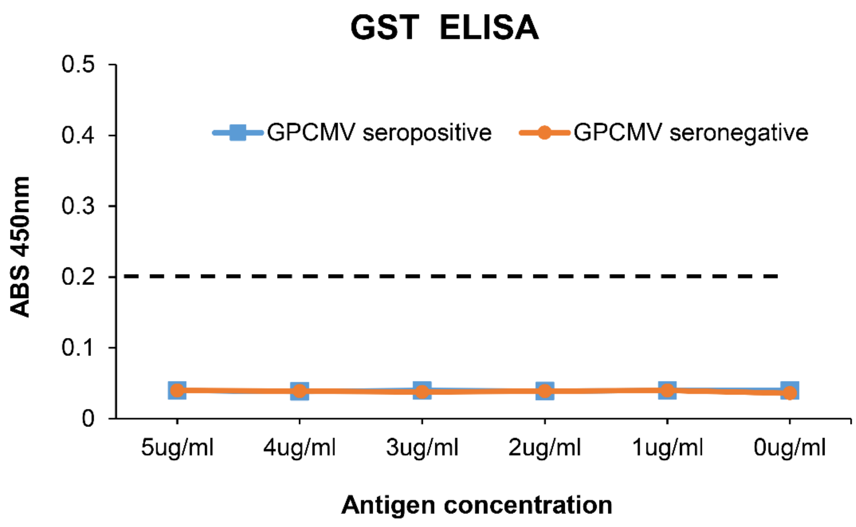


**Supplementary Figure 4. Antibody immune response to control GST antigen for GPCMV seropositive and seronegative sera.** Varying concentration of control carrier recombinant GST protein was used as coating antigen (0ug/ml to 5ug/ml) to test the reactivity of GPCMV seropositive and seronegative sera to GST. Assay negative cutoff for standard GPCMV ELISA assay, ABS 450nm $\leq$ 0.2, black dash line.
